# Plantar flexor muscle size, composition and strength and the redistribution of joint work in gait of healthy older adults

**DOI:** 10.64898/2026.01.30.702873

**Authors:** Ryan Gladfelter, Jane A Kent, Athulya A Simon, Katherine A Boyer

## Abstract

**Background:** Older adults exhibit hallmark changes in gait mechanics that may contribute to mobility loss. These changes are suggested to be precipitated by decreased mechanical output from the ankle plantar flexor muscles. However, the extent of age-related changes within individual triceps surae muscles and the consequences for plantar flexor specific torque (torque per unit muscle) and walking mechanics remains unclear. The study aims were to quantify age-related differences in triceps surae muscle cross-sectional area (CSA), fat fraction and peak absolute and specific plantar flexion torque, and to evaluate their relationships with positive hip and ankle work distribution during walking.

**Methods:** Eleven younger (36.2 +/-3.4 years, 5 female) and 12 older (73.3+/-2.8 years, 6 female) adults completed dynamometry, magnetic resonance fat-water imaging of the shank, and overground gait analysis at a prescribed (1.2 m/s) and fast walking speed.

**Results:** Older adults had a smaller fat-free muscle CSA for the soleus and medial gastrocnemius, resulting in ∽25% smaller triceps surae CSA. There was no difference by age in fat fraction for any muscle. Peak plantar flexor torque was lower in older and proportional to the differences in cross-sectional area, thus specific torque did not differ by age. Older adults generated greater positive hip work at both speeds and less positive ankle work at the fast speed leading to a greater redistribution ratio compared to younger. However, no significant correlations were found between the redistribution ratio and triceps surae muscle morphology or plantar flexion absolute or specific torque in older adults.

**Conclusions:** These results suggest that triceps surae muscle morphology and function may not be a primary source of age-related changes in gait mechanics in healthy older adults.

## Introduction

Older adults exhibit hallmark deficits in gait speed, step length, ankle kinetics, and propulsive force for reasons that are poorly understood (Boyer et al., 2012; Buddhadev et al., 2020; DeVita and Hortobagyi, 2000). Increased leading-limb hip extensor positive power during early stance may compensate for the decrease in ankle output (positive power and work) (Browne and Franz, 2019; Buddhadev et al., 2020; DeVita and Hortobagyi, 2000). This redistribution of positive joint power and work from the ankle to the hip has been referred to as the “Distal-to-Proximal Shift”. Some posit that the decrease in ankle mechanical output is an autonomous “decision” older adults make to avoid slips, trips, and falls (Quach et al., 2011; Sloot et al., 2021). However, evidence also suggests there may be limits in peak ankle moment production in older age (Browne and Franz, 2019; Conway and Franz, 2020), leading to lower ankle power generation and requiring proximal joint compensation to maintain walking speed. The consequences of shifting away from an ankle-centric gait pattern are not yet fully elucidated by it may disrupt normal stance-to-swing transitions and the forward acceleration of the body’s center (Zelik and Adamczyk, 2016) and may also be metabolically unfavorable (Delabastita et al., 2021; Umberger and Rubenson, 2011). Probing the factors that may contribute to limitations in ankle moment, power and work production, including triceps surae muscle morphology and function, may illuminate the causes for age-related changes in gait mechanics, mobility and the metabolic cost of walking (Boyer et al., 2023).

The triceps surae muscle group is responsible for generating ∽70-80% of the mechanical power necessary for ankle push-off and swing initiation, and is believed to be a primary modulator of gait speed and step length (Franz, 2016). These muscles experience age-related changes in muscle volume, neural activation and intramuscular fat (Hasson et al., 2011; Morse et al., 2004; Pinel et al., 2021; Winegard et al., 1996). Initial studies found the contractile volume of the triceps surae was ∽20% smaller (Hasson et al., 2011; Morse et al., 2005a) with a ∽4-fold greater intramuscular fat content in older compared to younger adults (Hasson et al., 2011; Pinel et al., 2021). The extent to which the individual triceps surae muscles atrophy in older age and contribute to lower plantar flexor peak torques is not well characterized. It has been suggested that there is greater atrophy in the gastrocnemii compared to the soleus with age, but limited data are available in mixed sex cohorts (Naruse et al., 2023). While the triceps surae muscles share a common tendon, there is evidence to support independent actions of the muscle sub-tendons and thus, there is potential for functional consequences of differential changes in muscles with age.

The mechanisms and consequences of intramuscular fat accumulation are not fully understood. *In silico* simulations suggest that intramuscular fat may resist fiber shortening, thereby impairing force production (Rahemi et al., 2015). The impact of greater intramuscular fat or changes in activation on muscle force production can be assessed in part by determining plantar flexor specific torque (torque produced per unit muscle contractile content). Studies of ankle plantar flexor specific torque are limited; however, initial work suggests older adults exhibit ∽20-30% lower specific torque values compared to younger adults(Csapo et al., 2014; Morse et al., 2004; Morse et al., 2005b). Regardless of its causes, lower triceps surae specific torque, indicative of impaired force production, may prompt, to a greater degree than atrophy alone, a redistribution of joint moments or power during gait. However, evidence to support a link between muscle specific torque and the distal-to-proximal shift is limited.

The extent of age-related changes in muscle size and fat content within individual triceps surae muscles and the consequences for plantar flexor specific torque and the distribution of joint power and work during gait remain unclear. Changes in lower extremity work distribution reflecting an age-related distal-to-proximal shift can be quantified using a redistribution ratio (Browne and Franz, 2018). Therefore, the first aim of this study was to quantify muscle fat-free CSA, volume, and fat fraction in the soleus, medial and lateral gastrocnemii muscles, as well as plantar flexor maximum voluntary contractions (MVC) and specific torque (Nm/CSA) in younger and older adults. We hypothesized that older adults would have smaller muscles and greater fat fractions in each muscle, and lower MVC and specific torque compared to younger adults. The second aim was to quantify differences between younger and older adults in gait kinetics, including peak ankle moment, propulsive ground reaction forces and the redistribution ratio, and examine the relationships between gait kinetics and muscle morphology. We hypothesized that older compared to younger adults would exhibit lower peak ankle moments and positive work and greater hip moments and positive work, resulting in a greater redistribution ratio and smaller propulsive ground reaction forces in the older group. Lastly, we hypothesized that individuals with larger redistribution ratios and smaller propulsive ground reactions forces would have smaller CSA, greater fat content and weaker plantar flexor muscles (maximum torque and specific torque).

## Materials and Methods

Eleven young (5 females) and 12 older (6 females) healthy adults free of any neurological or neuromuscular condition were studied (Table 1). The sample size was selected to ensure a power (β=0.8) to detect ∽ 30% difference between groups (α=0.05) in muscle volume and peak ankle moment using values from the literature (DeVita and Hortobagyi, 2000; Hasson et al., 2011). Study procedures were approved by the University of Massachusetts Amherst Institutional Review Board (protocol # 2040), and all participants provided written informed consent. Participants completed: plantar flexor MVCs on a dynamometer, overground gait analysis, magnetic resonance imaging of the shank, and one week of accelerometry to assess habitual physical activity levels.

**Table 1.** Group Characteristics. Group mean (SD), * indicates p≤0.05. PWS = preferred walking speed, F=female.

| **Table 1:** Group Characteristics. Group mean (SD), * indicates $p \leq 0.05$. PWS = preferred walking speed, F=female | | | | | | |
| --- | --- | --- | --- | --- | --- | --- |
| **Group** | **Age** | **Height (m)** | **Mass (kg)** | **PWS (m/s)** | **BMI (kg/m<sup>2</sup>)** | **MVPA (min/wk)** |
| **Younger** (n=11, 5F) | 36.2 (3.4) | 1.75 (8.2) | 86.3 (14.6) | 1.39 (0.2) | 28.0 (3.9) | 357 (186.7) |
| **Older** (n=12, 6F) | 73.3 (2.8) | 1.72 (8.8) | **70.2 (10.2)\*** | 1.38 (0.1) | **23.6 (3.1)\*** | 267 (113.3) |

### Physical Activity Measurements

To characterize our study groups, each participant wore a triaxial accelerometer (GT3X, ActiGraph, Pensacola, FL) over their right hip for 7-10 days. Weekly moderate-to-vigorous physical activity (MVPA) was calculated based on the first 7 days of valid use (10 hours of wear time) using established cut points (Troiano et al., 2008).

### Magnetic Resonance Imaging: Six-Point Dixon Technique

Six-point, fat-water imaging (2 stacks, 6-mm slice thickness, 80 slices, 1.25 x 1.25 x 6 mm voxel volume, echo times 2.46 and 6.15ms; 3.69 and 7.38ms;4.92 and 8.61ms) was used to obtain serial axial images of the shank musculature from the femoral condyles to the lateral malleolus. Images were obtained in a 3.0 T MRI scanner (3T Skyra, Siemens Medical Systems, Erlangen, Germany) using a 15-cm diameter extremity coil. Contiguous images were averaged to produce forty 12-mm slices along the full length of the shank (Gladfelter, 2024).

A custom *MATLAB* (Mathworks, Natick, MA) script was used to segment and analyze all images (Smith, 2013) (Figure 1). Segmentation began with the slice in which the soleus first appeared and ended with the slice immediately distal to the appearance of the femoral condyles. Shank length was calculated as the distance from the most lateral aspect of the lateral malleolus to the femoral condyles. For each muscle and slice, mean and SD fat percent (fat fraction, %FF) in the region of interest (ROI) were determined, and fat area was subtracted from total area to obtain the fat-free (i.e., muscle) area. Shank length, maximum fat-free muscle cross-sectional area (CSA_max_, cm^2^), and muscle volume (MV, cm^3^) were calculated. For each muscle, the overall FF (%) was calculated as the average FF for the 3 slices encompassing the CSA_max_. The CSA_max_ of the triceps surae (TS) muscle group was calculated as the sum of CSA_max_ for all individual muscles, regardless of location.

**Figure 1:**
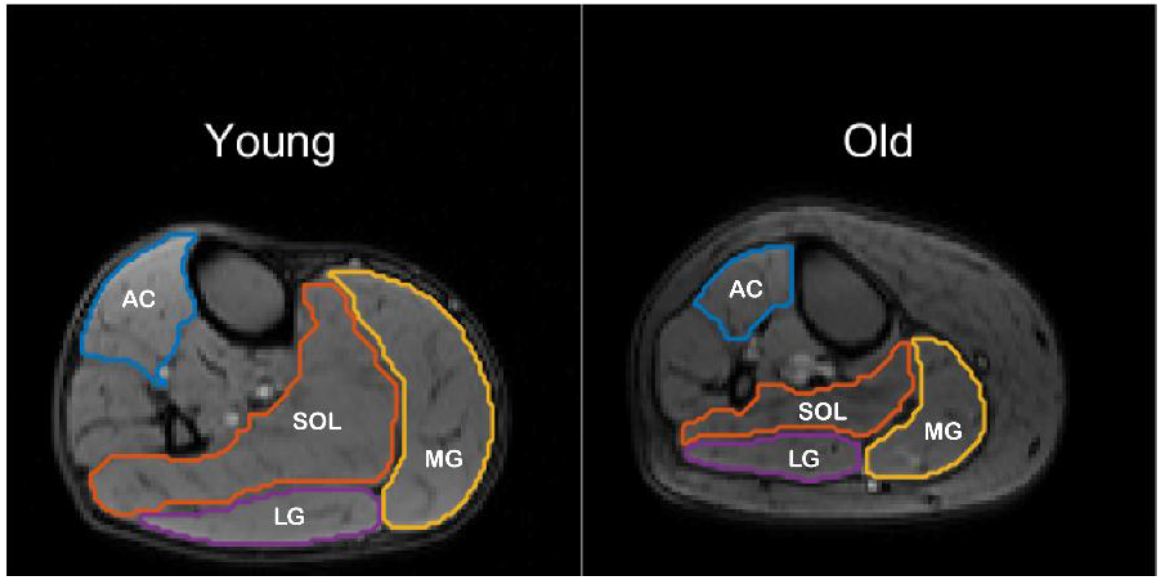
Representative muscle boundaries for single MRI slices from one younger (*left*) and one older (*right*) participant. Anterior compartment (Blue), soleus (Orange), medial (Yellow) and lateral (Purple) gastrocnemius regions of interest (ROI) are shown.

### Torque Measurements

Participants were seated on a Biodex dynamometer (Biodex System 4 Pro, NY) with the ankle at 90 degrees and knee fully extended, and completed three ankle plantar flexion MVCs (3-5s each, separated by 1min rest). Participants were instructed to “push as fast and hard as you can, like you are pressing down on a gas pedal”, and were given verbal encouragement during each contraction. Torque data was filtered with a 10 Hz low pass Butterworth filter and the maximum torque obtained over the 3 contractions was used for analysis. Ankle plantar flexor specific torque was calculated as peak torque divided by triceps surae muscle CSA_max_ (Nm.cm^-2^).

### Overground Gait Analysis

For overground gait analysis, seventy-three retro-reflective markers were placed bilaterally on bony landmarks as described previously (Boyer and Andriacchi, 2016). These markers were used to define the segment coordinate systems based on the ISB recommendations (Wu et al., 2002). Clusters of 9 and 6 markers were placed on each thigh and shank, respectively, to track the segment orientations during walking trials with the point cluster technique (Andriacchi et al., 1998; Boyer and Andriacchi, 2016). Following a standing trial, participants were instructed to walk across three in-ground force plates (AMTI Watertown, MA) while a 9 infrared camera system (Qualisys Oqus 7, Gothenburg, SE) tracked marker trajectories; 5 successful trials were averaged for each participant at each speed. Participants walked at a fixed speed of 1.2 m/s to minimize the effects of gait speed on our outcomes. Participants also walked at a speed that was 30 % faster than their preferred walking speed to test a challenging walking condition. Preferred walking speed was estimated as the average of 3 trials of a timed 6m walk, with a 3m rolling start (Johnson et al., 2020).

Visual3D with a custom *MATLAB* script was used to calculate joint kinematics and kinetics for the stance phase using an external, inverse dynamics approach (Andriacchi, 2004; Boyer and Andriacchi, 2016). Marker and force data were low-pass filtered at 8 and 15 Hz, respectively. The hip joint center was defined as in Bell (1990). Segment masses were defined using values provided by Dempster (1955) and the inertial parameters were defined using values from Hanavan (1964). Joint moments were resolved onto the distal segment and are reported as external moments. Joint powers were calculated as the product of joint moments and angular velocities. Positive mechanical joint work was calculated by integrating the positive joint power data with respect to time using the trapezium method.

The primary gait outcome measures were peak propulsive force, and peak ankle and hip moments, positive powers, and work. Secondary outcome measures included heel strike and peak joint angles and range of motion, and knee moments, powers, and work. External forces were scaled to body weight and joint moments were scaled to both bodyweight (N) and height (m), BWxHt. A joint work redistribution ratio was calculated as described previously (Browne and Franz, 2018). This redistribution ratio is bounded between 0 and 2, with 0 signifying all total positive work is done by the ankle, and 2 signifying all total positive work is done by the hip.

### Statistics

Shapiro-Wilks tests for normality were performed. When data were normally distributed, independent, one-sided t-tests were used to test for age-group differences; if not then non-parametric Mann-Whitney U tests were used (α=0.05). Pearson’s correlations (r) or non-parametric Spearman Rank (rs) tests were used to evaluate correlations between both redistribution ratio and propulsive force and: MVC, specific torque, triceps surae CSA_max,_ and fat fraction for all participants and by age group. Statistics were conducted using IBM SPSS Statistics v.29.0.1.0. Means, SD, effect sizes (*d*) and precise p values are reported.

## Results

The groups did not differ by height, but younger had greater mass and BMI than older (p ≤ 0.004), Table 1. There was no difference in preferred walking speed (p = 0.8) or weekly MVPA (p=0.09).

### Muscle Size

Triceps surae and medial gastrocnemius CSA_max_ (U= 106.0, p=0.01, *d=*1.2; U = 109.0, p=0.007, *d*=1.35) and MV (t(21) = 2.28, p=0.02, *d*=0.95; U=109.0, p = 0.009, *d*=1.47) were smaller in older than younger adults (Figure 2); as was soleus CSA_max_ (t(df)= 2.46, p=0.01, *d*=1.03; Table 2). The differences were not significant for the lateral gastrocnemius (CSA U=96.5, p = 0.06, *d*=0.97, MV U=82.0, p=0.35, d=0.62) or soleus MV (U=89.0. p=0.16, *d*=0.70). There were no effects of age on FF in any muscle (p≥0.34). Shank length and the percentage of shank length analyzed for each muscle (Table 2) did not differ by group.

**Figure 2:**
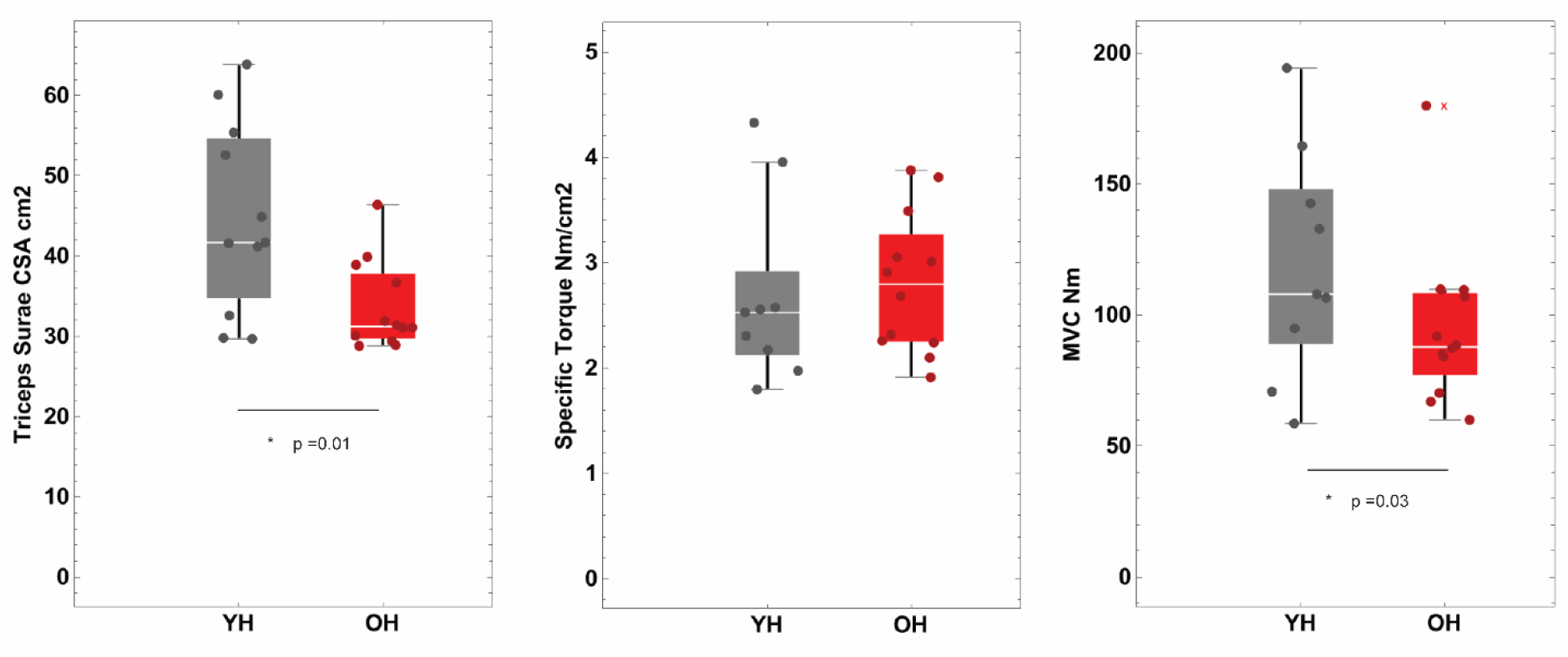
Triceps surae muscle size, torque and specific torque. The triceps surae muscle was smaller in older than younger adults (*left*), as was maximal torque (*middle*). Muscle specific torque was not different by group (*right*); x indicates outlier excluded from analysis.

**Table 2.**
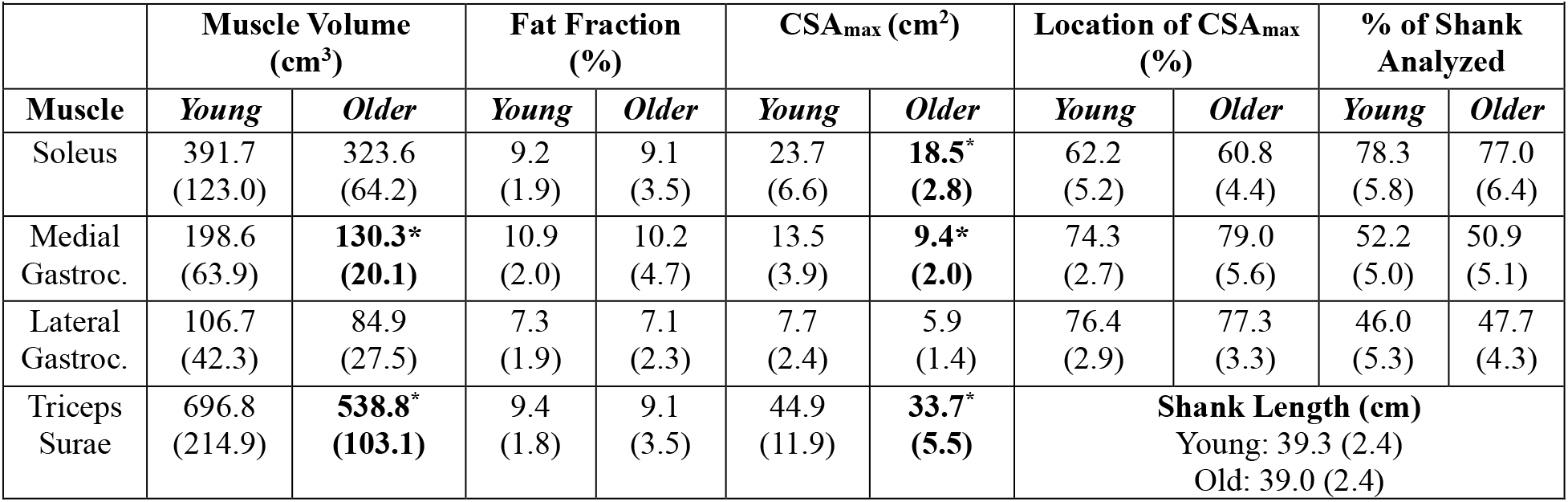
MRI measures. Values are Group mean (SD) * p≤0.05, Location of CSA_max_ (%): 0% represents lateral malleoli, 100% represents femoral condyles.

|  | Muscle Volume (cm<sup>3</sup>) |  | Fat Fraction (%) |  | CSA<sub>max</sub> (cm<sup>2</sup>) |  | Location of CSA<sub>max</sub> (%) |  | % of Shank Analyzed |  |
| --- | --- | --- | --- | --- | --- | --- | --- | --- | --- | --- |
| Muscle | Young | Older | Young | Older | Young | Older | Young | Older | Young | Older |
| Soleus | 391.7 (123.0) | 323.6 (64.2) | 9.2 (1.9) | 9.1 (3.5) | 23.7 (6.6) | **18.5\* (2.8)** | 62.2 (5.2) | 60.8 (4.4) | 78.3 (5.8) | 77.0 (6.4) |
| Medial Gastroc. | 198.6 (63.9) | **130.3\* (20.1)** | 10.9 (2.0) | 10.2 (4.7) | 13.5 (3.9) | **9.4\* (2.0)** | 74.3 (2.7) | 79.0 (5.6) | 52.2 (5.0) | 50.9 (5.1) |
| Lateral Gastroc. | 106.7 (42.3) | 84.9 (27.5) | 7.3 (1.9) | 7.1 (2.3) | 7.7 (2.4) | 5.9 (1.4) | 76.4 (2.9) | 77.3 (3.3) | 46.0 (5.3) | 47.7 (4.3) |
| Triceps Surae | 696.8 (214.9) | **538.8\* (103.1)** | 9.4 (1.8) | 9.1 (3.5) | 44.9 (11.9) | **33.7\* (5.5)** | **Shank Length (cm)**<br>Young: 39.3 (2.4)<br>Old: 39.0 (2.4) |  |  |  |

### Muscle Torque

Plantar flexor peak and specific torques are shown in Figure 2. Plantar flexor MVC was 27% lower in older than younger (t(18) = -2.23, p=0.02, *d*=0.65). Specific torque (U=62.0, p=0.60, *d*=0.15) was not different between groups. One younger female and one younger male were excluded due to missing data.

### Gait – Prescribed Speed

At the prescribed walking speed, all participants were able to walk within 5% of the 1.25 m/s and thus walking speed was not different between groups (Table 3).

**Table 3.** Discrete joint kinematic and kinetic values for older and younger groups at the fixed walking speed of 1.2m/s. Values are group means (SD). * p≤0.05 Precise p-value and cohen’s d effects sizes are reported. ^+^ indicates Mann -Whitney U test for group differences.

| **Table 3:** Discrete joint kinematic and kinetic values for older and younger groups at the fixed walking speed of 1.2m/s. Values are group means (SD). * $p \leq 0.05$ Precise p-value and cohen's d effects sizes are reported. + indicates Mann -Whitney U test for group differences. | | | | |
| --- | --- | --- | --- | --- |
| **ANKLE** | **Young** | **Old** | **p-value** | **Effect Size** |
| Gait Speed (m/s) | 1.25 (0.05) | 1.24 (0.02) | 0.18 | 0.40 |
| HS Ankle Flexion (deg.) | 3.97 (3.18) | 2.52 (3.79) | 0.17 | 0.41 |
| **Peak Ankle Plantar Flexion (deg.)\*** | **-18.21 (4.23)** | **-14.37 (5.54)** | **0.04\*** | **0.76** |
| Stance Ankle Range of Motion (deg.) | 30.29 (4.16) | 29.17 (4.48) | 0.28 | 0.26 |
| Peak Ankle Plantar Flexion Moment (%BW *HT) | 1.38 (0.39) | 1.68 (0.60) | 0.10 | 0.57 |
| Peak Ankle Dorsiflexion Moment (%BW *HT) | -8.17 (0.97) | -8.02 (0.80) | 0.34 | 0.18 |
| Peak Positive Ankle Power (W/kg) | 2.74 (0.49) | 2.55 (0.41) | 0.17 | 0.41 |
| Total Positive Ankle Work (J/kg) | 0.25 (0.05) | 0.22 (0.05) | 0.12 | 0.52 |
| **KNEE** |  |  |  |  |
| **HS Knee Flexion (deg.)\*** | **-0.47 (6.61)** | **4.62 (5.22)** | **0.03\*** | **0.81** |
| **Mid-Stance Knee Flexion (deg.)\*** | **15.55 (6.57)** | **22.46 (6.19)** | **0.01\*** | **0.97** |
| Knee Flexion Range of Motion (stance) (deg.) | 46.00 (4.10) | 44.09 (4.30) | 0.15 | 0.16 |
| Peak Knee Extension Moment (%BW *HT) | -2.19 (0.91) | -2.11 (1.17) | 0.43 | 0.16 |
| Peak Knee Flexion Moment (%BW *HT) | 2.87 (0.10) | 3.04 (1.20) | 0.36 | 0.08 |
| Peak Positive Knee Power (W/kg) | 0.67 (0.26) | 0.67 (0.26) | 0.49 | 0.01 |
| Total Positive Knee Work (J/kg) | 0.11 (0.04) | 0.10 (0.04) | 0.30 | 0.23 |
| **HIP** |  |  |  |  |
| Hip Range of Motion (stance) (deg.) | 39.96 (6.57) | 40.86 (7.15) | 0.38 | 0.13 |
| **Peak Hip Flexion Moment (%BW \*HT)** | **-2.50 (0.50)** | **-3.85 (1.67)** | **0.01\*** | **0.95** |
| Peak Hip Extension Moment (%BW *HT) | 5.45 (1.86) | 4.61 (1.49) | 0.13 | 0.50 |
| +Peak Positive Hip Power (W/kg) | 1.40 (0.40) | 1.73 (0.67) | 0.18 | 0.55 |
| **Positive Hip Work \*(J/kg)** | **0.10 (0.02)** | **0.14 (0.08)** | **0.04\*** | **0.78** |
| **LIMB** |  |  |  |  |
| Peak Propulsive force (%BW) | 0.20 (0.01) | 0.19 (0.02) | 0.07 | 0.65 |
| Total Limb Positive Work (J/kg) | 0.46 (0.06) | 0.47 (0.08) | 0.34 | 0.19 |
| **Redistribution Ratio** | **0.56 (0.15)** | **0.76 (0.27)** | **0.03** | **0.87** |

Older adults walked with smaller peak plantar flexion angles (t(20) = -1.9, p=0.04) but the ankle angle at heel-strike and range of motion were not different from young (t(20)-.96, p=0.17, t (20) = 0.60, p=0.28). No group differences in the peak ankle moments (t(20) = -1.35, p=0.1; t(20) =0.41, p=0.34) positive power (t(20) = 0.97, p=0.17) or work (t(20) =1.2, p=0.12) were found (Table 3, Figure 3).

**Figure 3:**
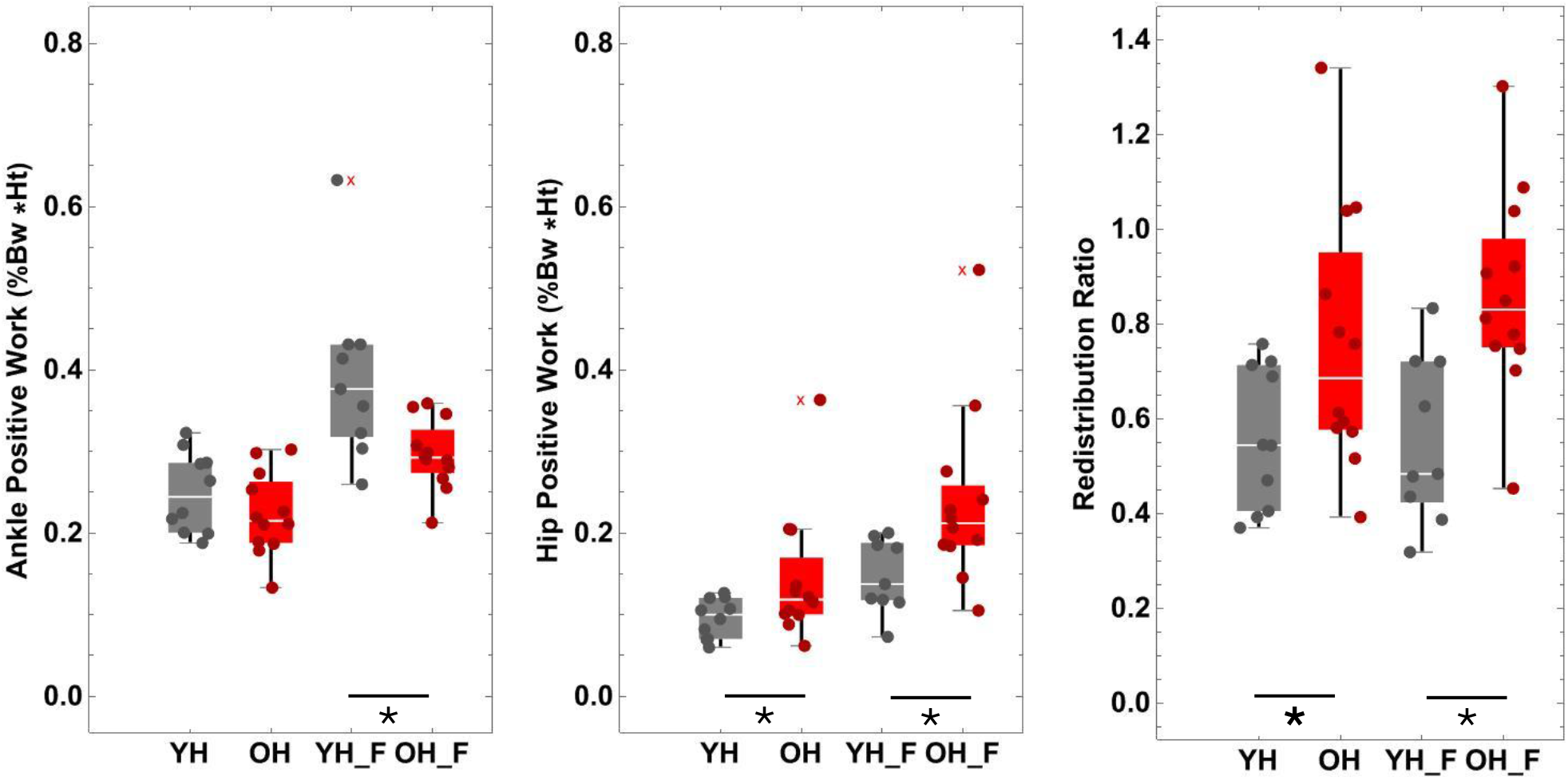
Positive ankle work (*left*), hip positive work (*middle*) and redistribution ratio (*right*), for each group at both prescribed and fast walking speeds. The redistribution ratio was greater in older due to greater hip positive work at both speeds as well as smaller ankle positive work for fast walking. No group differences were found for ankle positive work. Outlier (x) were not excluded as are plausible values based on larger datasets

Older adults had greater knee flexion at heel-strike (t(20) = -2.0, p=0.03) and in midstance (t(20)=-2.5, p=0.01) but no differences were found in the stance phase knee range of motion (t(20)= 1.1, p=0.15). There were no age group differences in the knee moments, positive power or work (Table 3).

No differences in hip range of motion were found between older and younger adults (t(20) = -0.30, p = 0.38). Older adults had greater hip flexion moments in early stance (t(20) =2.5, p=0.01) and positive hip work (t(20) = -1.8, p=0.04, Figure 3) but there were no differences in the hip extension moment (t(20) = 1.2, p=0.13) or peak positive hip power (U=81.0 p=0.18), Table 3.

The total limb positive work (t(20) =-0.4, p=0.34) and propulsive ground reaction force (t(20) = 1.5, p= 0.07) were not different in the older adults but the redistribution ratio was greater for older adults (t(20) =-2.0, p = 0.03), Table 3, Figure 3.

### Gait – Fast speed

For fast walking, data were not available for 2 younger participants due to occlusion of tracking markers. Fast walking speeds were ∽30% faster than participants’ preferred walking speeds and were also not different between groups (t(19)=0.68, p=0.25) (Table 4).

**Table 4.** Discrete ankle joint kinematic and kinetic values for older and younger **groups at faster than preferred** walking speed. Values are group means (SD). * p≤0.05 Precise p-value and cohen’s d effects sizes are reported. ^+^ indicates Mann -Whitney U test for group differences.

| ANKLE | Young | Old | p-value | Effect Size |
| --- | --- | --- | --- | --- |
| Gait Speed (m/s) | 1.88 (0.25) | 1.82 (0.1) | 0.25 | 0.30 |
| HS Ankle Flexion (deg.) | 8.43 (5.2) | 6.70 (4.1) | 0.20 | 0.38 |
| **Peak Ankle Plantar Flexion (deg.)\*** | **-23.66 (5.7)** | **-16.99 (6.0)** | **0.009** | **1.13** |
| **Stance Ankle Range of Motion (deg.)** | **35.96 (7.23)** | **29.49 (4.38)** | **0.01** | **1.12** |
| +Peak Ankle Plantar Flexion Moment (%BW *HT) | 2.16 (0.76) | 2.57 (0.0.96) | 0.55 | 0.47 |
| Peak Ankle Dorsiflexion Moment (%BW *HT) | -9.28 (1.7) | -8.99 (0.9) | 0.34 | 0.23 |
| +Peak Positive Ankle Power (W/kg) | 4.38 (1.48) | 3.88 (0.7) | 0.46 | 0.51 |
| **Total Positive Ankle Work J/kg)** | **0.39 (0.11)** | **0.30 (0.04)** | **0.006** | **1.24** |
| KNEE |  |  |  |  |
| **HS Knee Flexion (deg.)\*** | **0.3 (7.44)** | **7.59(5.68)** | **0.01** | **1.12** |
| Mid-Stance Knee Flexion (deg.)* | 20.92 (10.02) | 26.44 (5.89) | 0.06 | 0.70 |
| Knee Flexion Range of Motion (stance) (deg.) | 41.38 (5.82) | 42.47 (3.43) | 0.29 | 0.24 |
| Peak Knee Extension Moment (%BW *HT) | 2.87 (1.17) | 2.93 (1.04) | 0.45 | 0.05 |
| Peak Knee Flexion Moment (%BW *HT) | 5.28 (2.24) | 5.44 (1.67) | 0.43 | 0.08 |
| Peak Positive Knee Power W/kg) | 1.19 (0.31) | 1.26 (0.49) | 0.37 | 0.15 |
| Total Positive Knee Work J/kg) | 0.20 (0.07) | 0.18 (0.07) | 0.31 | 0.22 |
| HIP |  |  |  |  |
| Hip Range of Motion (stance) (deg.) | 45.83(7.45) | 49.29 (5.79) | 0.12 | 0.52 |
| **Peak Hip Flexion Moment (%BW \*HT)** | **-5.33 (1.38)** | **-7.30 (2.75)** | **0.03** | **0.81** |
| Peak Hip Extension Moment (%BW *HT) | 7.23 (1.90) | 6.74 (2.00) | 0.29 | 0.25 |
| +Peak Positive Hip Power W/kg) | 2.66 (0.83) | 3.59 (1.10) | 0.06 | 0.86 |
| **+Positive Hip Work \*(J/kg)** | **0.15 (0.04)** | **0.24 (0.11)** | **0.01** | **0.93** |
| Peak Propulsive force (%BW) | 0.28 (0.06) | 0.25 (0.03) | 0.04 | 0.76 |
| Total Limb Positive Work (J/kg) | 0.74 (0.13) | 0.72 (0.14) | 0.37 | 0.15 |
| **Redistribution Ratio\*** | **0.56 (0.18)** | **0.86 (0.22)** | **0.001** | **1.23** |

At the fast speed, older adults walked with a smaller peak ankle plantarflexion angle (t(19)=-2.6, p=0.009) and angle range of motion(t(19)=2.6, p=0.01) (Table 4). While the ankle moments and peak positive power were not different between groups at the fast speed (U=63.0, t(19)=-0.52, U=43.0, p>0.05), ankle positive work was 23% lower for older as compared to young (t(19)= 2.8, p=0.006) (Figure 3).

Older adults walked with greater knee flexion at both heel strike (t(df)= -2.6, p=0.01) and mid -stance (t(19)=-1.6, p=0.06) but the total knee range of motion in stance was not different between older and young (t(19)=-0.54, p=0.17). There were no age group differences in the knee moments, positive power or work in fast walking (t(19)=-0.19, t(19)= 0.12, t(19)= -0.34, t(19)=0.49, p>0.05) (Table 4).

No differences were found in hip range of motion between age groups (t(df)=-1.2, p=0.12). Older adults had a greater hip flexion (t(19)=1.96, p=0.03) but not extension moment (t(19)=0.56, p=0.29), slightly greater hip positive power (U=81.0,p=0.06) and greater hip positive work (U=89.0, p=0.01) for fast walking.

There was no difference in total limb work between age groups (t(19)=0.34, p=0.37). The peak propulsive force was smaller (t(19)=1.8, p=0.04) and there was a larger redistribution ratio (t(df)=-3.5, p=0.001) for older adults as compared to younger adults.

### MRI, Torque, and Gait Relationships

At the prescribed speed, there were no correlations between the redistribution ratio and: plantar flexor specific torque (r_s_=0.28, p=0.21), MVC (r_s_=-0.09, p=0.70), triceps surae CSA_max_ (r_s_=-0.33, p=0.13) or triceps surae fat fraction (r=0.03, p=0.89). Similarly, there were no correlations between propulsive force: and plantar flexor specific torque (r_s_=0.02, p=0.91), MVC (r_s_=0.35, p=0.12), or triceps surae CSA_max_ (r_s_=0.28, p=0.20). There was a negative correlation between propulsive force and triceps surae fat fraction (r_s_=-0.44, p=0.04) for the groups combined. When the groups were considered separately, this association was significant only for the older group (older r_s_=-0.56, p=0.05; younger r_s_=-0.36, p=0.30).

At the fast speed, there was a significant correlation between redistribution ratio and triceps surae CSA_max_ (r_s_=-0.54,p=0.01) for the groups combined. When the groups were considered separately, this association was significant only for the younger group (older r_s_ = 0.15,p=0.64; younger: r_s_=-0.73,p=0.02), Figure 4. The redistribution ratio was not correlated with MVC (r_s_=-0.34, p=0.15, Figure 4), fat fraction (r=-0.006, p=0.98), or specific torque (r=0.06, p=0.79). There was a negative correlation between propulsive force and triceps surae fat fraction (r=-0.47, p=0.03) for the groups combined. When the groups were considered separately, this association was significant only for the older group (older r=-0.69, p=0.01; younger r=-0.43, p=0.24). The propulsive force was also correlated with MVC (r_s_=0.51, p=0.02) when groups were combined but was not significant for either group when tested alone (older r=0.22, p =0.62; younger r=0.27, p=0.38). There were no correlations between propulsive force and triceps surae CSA_max_ (rs=0.2,p=0.36) or specific torque (r=0.16, p=0.50).

**Figure 4.**
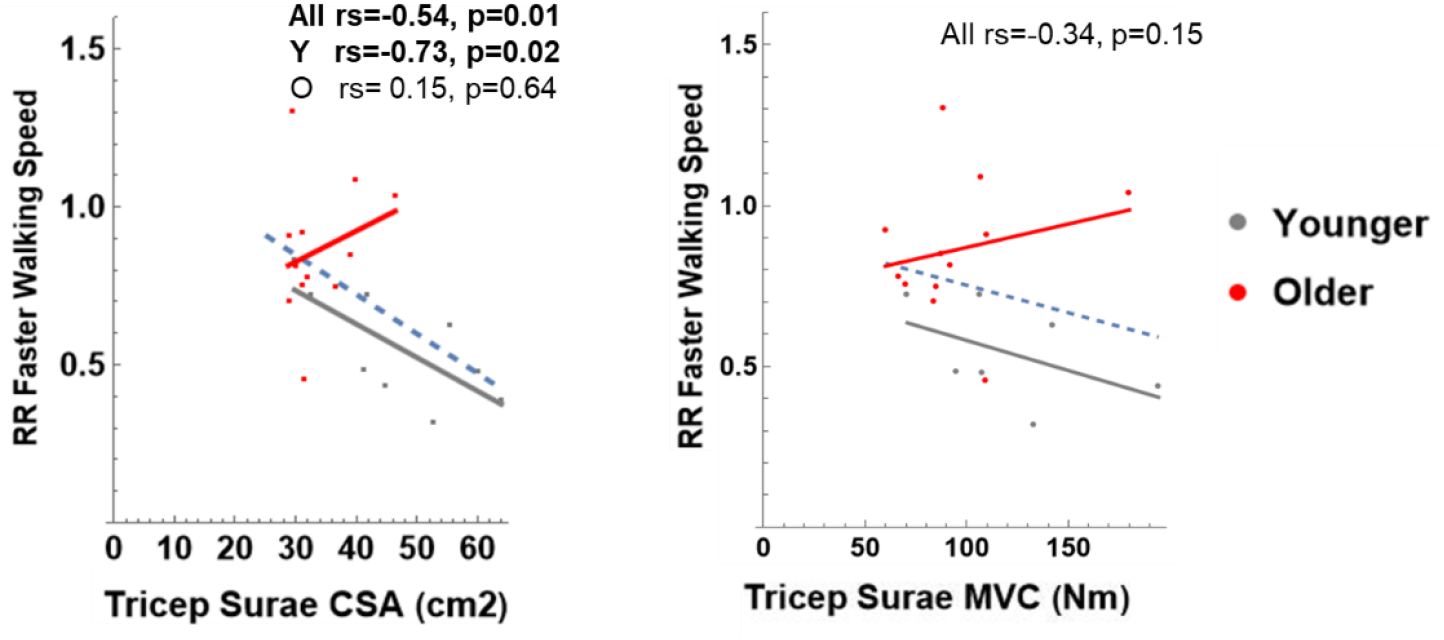
While there was a significant correlation between the redistribution ratio at the fast walking speed and triceps surae CSA (dashed line, right panel), this relationship was only significant for the younger group (gray solid line) when age groups were tested separately. There was not a correlation between the redistribution ratio and triceps surae MVC (left panel).

## Discussion

Our aim was to quantify age-related differences in triceps surae muscle morphology and to evaluate the potential consequences for plantar flexor absolute and specific torque and walking mechanics. Older adults had smaller soleus, medial gastrocnemius and triceps surae CSAs, with no difference by age in fat fraction for any muscle. Isometric plantar flexor strength (MVC) was lower in older by approximately the same magnitude as triceps surae size; thus, specific torque did not differ by age. When walking at 1.2m/s, older adults generated greater total positive hip work and had a greater redistribution ratio compared with younger adults. At 30% faster than their preferred walking speed, older adults generated greater hip positive work as well as less ankle positive work than younger adults and thus age differences in the redistribution ratio were increased. However, at neither the prescribed nor faster speed were the redistribution ratio or propulsive ground reaction forces during gait associated with triceps surae muscle size or function in older adults. Together, these results suggest that deficits in triceps surae size and isometric strength do not impair gait function in healthy older adults and thus are unlikely to be essential factors in the age-related redistribution of joint work in walking.

There are conflicting reports regarding age-related differences in triceps surae muscle size. We observed an ∽23-25% smaller volume and CSA_max_ in triceps surae of older compared with younger adults, which is consistent with the ∽20% smaller volume for older adults reported in the literature (Barber et al., 2013; Csapo et al., 2014; Hasson et al., 2011; Morse et al., 2005a). The literature indicates that older adults display smaller muscle volumes in each muscle of the triceps surae(Morse et al., 2005a), in the gastrocnemii only (Barber et al., 2013), or in none of the muscles (Csapo et al., 2014; Pinel et al., 2021; Zange et al., 2025). We found the largest age-related difference in size to be in the medial gastrocnemius. The lack of objective measures of physical activity in prior work makes it difficult to interpret the conflicting findings in the literature. Our findings in a population of older adults who on average met or exceeded the ACSM recommendations for aerobic exercise (> 150 MVPA/Week), extend the prevailing literature suggesting that the age-related decrease in triceps surae size (CSA_max_ and volume) is due to variable contributions across muscles. The atrophy of individual triceps surae muscles may be important to consider as, although they share a common tendon, these muscles may have different contributions to propulsion and support, and may have independent actuation abilities (Clark and Franz, 2020; Clark et al., 2021; Neptune et al., 2001).

Fat-free muscle CSA provides a good representation of the number of sarcomeres in parallel and thus should predict the capacity for isometric torque production. Peak isometric torque was lower in the older adults, as expected based on their smaller triceps surae CSA_max_. In contrast to our hypothesis, no differences in specific torque between younger and older adults were found, suggesting no inherent deficit in contractile function in the smaller muscles of the older group. There were also no age-related differences in fat fraction in any plantar flexor muscle, which is a possible contributor to lower specific strength in older. This finding conflicts with some literature, as others have reported greater intramuscular fat or non-contractile content in the triceps surae muscles of older adults (Csapo et al., 2014; Hasson et al., 2011; Pinel et al., 2021; Schwenzer et al., 2009; Zange et al., 2025). The fat fraction in our older group is similar to what has been reported previously using a similar fat-water imaging sequence (Zange et al.,2025), so the lack of age-group differences may come from a greater fat content in our younger group. There is growing evidence that age-related differences in torque production and thus specific torque may be greater for dynamic compared with isometric contractions (Callahan and Kent-Braun, 2011; Fitzgerald et al., 2021; Thompson et al., 2013). The consequences of changes in triceps surae muscle morphology on specific dynamic torque remain unclear.

In agreement with our hypothesis, older adults had a greater redistribution ratio compared to young adults, indicating a greater reliance on hip musculature for positive work generation during both prescribed and faster speed walking. At the prescribed speed, the peak external hip flexion moment and positive hip work were both greater in older, compared to younger, adults. However, neither positive ankle power or work, nor the peak external ankle dorsiflexion moment were different between age groups at the prescribed speed. As a result, the redistribution ratio, while larger for older adults, was smaller than that previously reported for older adults in the literature (Browne and Franz, 2018). Prior work has suggested the redistribution of lower limb joint work and smaller propulsive force is due to ankle power deficits in late stance or limitations in ankle moment production that are compensated for by leading limb hip extensor power during early stance (Browne and Franz, 2019; Buddhadev et al., 2020; Conway and Franz, 2020; DeVita and Hortobagyi, 2000). The lack of differences in ankle kinetics suggests that ankle moment production was not limited in the older adults. The prescribed speed of 1.2 m/s was ∽10% slower on average than the preferred walking speeds of the study participants. The lesser demands of this task may have been well within the capacity of the lower limb muscles to produce the required joint moments and powers.

Age related gait differences were more evident at the faster walking speed. At the fast speed, older adults walked with a smaller ankle range of motion and less positive ankle work, but greater hip flexion moments and hip positive work leading to a larger redistribution ratio than young. For younger adults the mean redistribution ratio was not different between walking speeds. However, for older adults, the redistribution ratio was ∽13% greater at the fast compared with prescribed speed, suggesting that the greater task demand inherent in fast walking required increased relative effort from weakened triceps surae muscle groups, eliciting an altered gait pattern. The faster speed was set at 30% faster than individuals’ preferred walking speed and approached the walk to run transition speed for most participants (Farinatti and Monteiro, 2010) suggesting it was likely near each person’s maximum walking speed. Older adults had weaker and smaller plantar flexor muscles; however, the peak ankle moment was not smaller in older adults, suggesting limitations in ankle moment production may not be a key factor in the changes in propulsive forces and the redistribution ratio(Conway and Franz, 2020). Although not significant, the peak positive ankle power in walking was 11% smaller in older adults contributing to the lower ankle positive work. Some studies suggests that deficits in muscle power production may be a stronger and earlier predictor of functional decline in older adults than isometric torque (Bean et al., 2003; Reid and Fielding, 2012). Although the plantar flexor isometric peak torque (MVC) was smaller in older, the lack of differences in the peak ankle moment but decreased ankle power production during gait suggest that future studies should include assessments of both isometric and dynamic plantar flexor torque production to further investigate age-related alterations in muscle torque and power production impact gait.

To probe how variation in triceps surae muscle morphology or function may impact gait, we tested for correlations between the redistribution ratio and propulsive force with cross sectional area, fat fraction, and absolute and specific plantar flexion torque. In fast walking, a significant relationship between the redistribution ratio and triceps surae CSA was found when all participants were grouped together, however when separated by age a significant relationship was only found for young. There were no correlations between the redistribution ratio or propulsive ground reaction forces and any of the function measures. The lack of a relationship between redistribution ratio and absolute or specific torque, although surprising, is supported by some prior work that showed strengthening the triceps surae has no effect on habitual ankle kinetics or preferred walking speed (Beijersbergen et al., 2013; Conway and Franz, 2020). It may be that factors contributing to the change in gait mechanics with age might be more systemic (e.g. proximal or whole limb muscle function) or complex (e.g. multi-joint compensation) than simply small deficits in plantar flexor muscles moment generation. There was a correlation between the fat fraction and propulsive force, yet no difference in fat fraction by age group. The mechanisms and consequences of intramuscular fat are still not yet fully understood. Evidence from *in silico* simulations suggests that fat increases the base stiffness of muscle, and this increase in stiffness would limit muscle fiber shortening and axial bulging (Rahemi et al., 2015), thus impacting force production. Given the fat fraction was not different by age group, it is not possible to speculate on the impact of intramuscular fat on lower limb work distribution with age.

We did not incorporate calcaneal tendon measures in this study, although older adults generally have more compliant tendons with larger cross-sectional areas and smaller moment arms compared to younger adults (Epro et al., 2018; Krupenevich et al., 2022; Rasske and Franz, 2018; Stenroth et al., 2012). Future mechanistic studies that evaluate the whole muscle-tendon unit, as well additional factors such as muscle architecture (e.g., fascicle length, pennation angle), neural activation, and intramuscular fat content will advance our understanding of gait-related changes in older age. Finally, there remains a need to quantify the influence of triceps surae muscle morphology and function on gait mechanics in older adults with a range of physical activity levels, as those in this study were all moderately active.

In conclusion, the results of this study provide support for muscle-specific changes within the triceps surae in older age, with the greatest age-related changes in muscle size occurring in the medial gastrocnemius. Notably, we did not find greater intramuscular fat or lower specific torque in these healthy older adults, nor were there direct implications for gait mechanics of the smaller and weaker triceps surae muscle group for the older adult group. Older adults did have greater reliance on the hip than younger adults, reflected by their greater redistribution ratio at both preferred and fast walking speeds. However, there was no relationship of the redistribution ratio with triceps surae muscle size or function in older adults. Together, these results suggest that triceps surae muscle morphology and function may not be essential factors in the age-related changes in gait mechanics for healthy, physically-active older adults.

## Acknowledgements

We thank the participants for their time, Amir Shah for assistance in MRI analysis, Skylar Holmes PhD for assistance in data collection and Zoe Smith, MS and Jacob Thomas, PhD for their insightful comments in the preparation of this manuscript.

